# Structural Genetics of circulating variants affecting the SARS-CoV-2 Spike / human ACE2 complex

**DOI:** 10.1101/2020.09.09.289074

**Authors:** Francesco Ortuso, Daniele Mercatelli, Pietro Hiram Guzzi, Federico Manuel Giorgi

## Abstract

SARS-CoV-2 entry in human cells is mediated by the interaction between the viral Spike protein and the human ACE2 receptor. This mechanism evolved from the ancestor bat coronavirus and is currently one of the main targets for antiviral strategies. However, there currently exist several Spike protein variants in the SARS-CoV-2 population as the result of mutations, and it is unclear if these variants may exert a specific effect on the affinity with ACE2 which, in turn, is also characterized by multiple alleles in the human population. In the current study, the GBPM analysis, originally developed for highlighting host-guest interaction features, has been applied to define the key amino acids responsible for the Spike/ACE2 molecular recognition, using four different crystallographic structures. Then, we intersected these structural results with the current mutational status, based on more than 295,000 sequenced cases, in the SARS-CoV-2 population. We identified several Spike mutations interacting with ACE2 and mutated in at least 20 distinct patients: S477N, N439K, N501Y, Y453F, E484K, K417N, S477I and G476S. Among these, mutation N501Y in particular is one of the events characterizing SARS-CoV-2 lineage B.1.1.7, which has recently risen in frequency in Europe. We also identified five ACE2 rare variants that may affect interaction with Spike and susceptibility to infection: S19P, E37K, M82I, E329G and G352V.

**Significance Statement:** We developed a method to identify key amino acids responsible for the initial interaction between SARS-CoV-2 (the COVID-19 virus) and human cells, through the analysis of Spike/ACE2 complexes. We further identified which of these amino acids show variants in the viral and human populations. Our results will facilitate scientists and clinicians alike in identifying the possible role of present and future Spike and ACE2 sequence variants in cell entry and general susceptibility to infection.

## Main Text

## Introduction

The Severe Acute Respiratory Syndrome Coronavirus 2 (SARS-CoV-2) has emerged in late 2019 (1) as the etiological cause of a pandemic of severe proportions dubbed Coronavirus Disease 19 (COVID-19). The disease has reached virtually every country in the globe (2), with more than 40,000,000 confirmed cases and more than 1,100,000 deaths (source: World Health Organization). SARS-CoV-2 is characterized by a 29,903-long single stranded RNA genome, densely packed in 11 Open Reading Frames (ORFs); the ORF1 encodes for a polyprotein which is furtherly split in 16 proteins, for a total of 26 proteins (3).

The second ORF encodes for the Spike (S) protein, which is the key protagonist in the viral entry into host cells, through its interaction with human epithelial cell receptors Angiotensin Converting Enzyme 2 (ACE2) (4), Transmembrane Serine Protease 2 (TMPRSS2) (5), Furin (6) and CD147 (7). Investigators have focused their attention on the Spike/ACE2 interaction, trying to disrupt it as a potential anti-COVID-19 therapy, using small drugs (8) or Spike fragments (9). Using X-ray crystallography, some models of the Spike/ACE2 have been generated (10–12), providing a structural instrument for the analysis of this key interaction. These models determined that the Receptor Binding Domain (RBD) of Spike, directly interacting with ACE2, is a compact structure of ∼200 amino acids (AAs) over a total of 1273 AAs of the full-length Spike.

The SARS-CoV-2 Spike protein adapted from subsequent mutations from a wild bat beta-coronavirus (13), in order to exploit the N-terminal ACE2 peptidase domain conformation. As a result, SARS-CoV-2 Spike can establish a strong interaction with the human cell surface, allowing the virus to fuse its membrane with that of the host cell, releasing its proteins and genetic material and starting its replication cycle (5). While SARS-CoV-2 shows low mutability (14), with less than 25 predicted events/year (15), the virus is in continuous evolution from the original Wuhan reference sequence (NC_045512.2) (16), and there are currently at least 6 major variants circulating in the population (3, 17). Some of these strains are characterized by a mutation in Spike, at AA 614, whereas an Aspartic Acid (D) is substituted by a Glycine (G) (18). In fact, the Spike D614G mutation gives the name to the most frequent viral clade (G), which was first detected in Europe at the end of January 2020, and is currently present in all continents, with increasing frequency over time (3). D614G does not fall within the putative RBD (AA ∼330-530), but some studies suggest it may have a clinically relevant role: D614G is positively correlated with increased case fatality rate (19), and it shows increased transmissibility and infectivity compared to the reference genome (20). *In vitro* studies show that viruses carrying the D614G Spike mutation have an increased viral load and cytopathic effect in cultured Vero cells (16). Despite these preliminary observations, there are still several doubts on the molecular effects of the D614G variant (21). Other recurring Spike mutations have been observed in the population worldwide, however at frequencies of 1% or below (3); some of these mutations fall within the RBD and therefore may have a direct role in ACE2 interaction.

On the other hand, genetic variants of ACE2 in human population may influence susceptibility or resistance to SARS-CoV-2 infection, possibly contributing to the difference in clinical features observed in COVID-19 patients (22). ACE2 gene is located on chromosome Xp22.2 and consists of 18 exons, coding for an 805 AAs long protein exposed on the cell surface of a variety of human organs, including kidneys, heart, brain, gastrointestinal tract, and lungs (23). It is unclear if tissue-expression patterns of ACE2 may be linked to the severity of symptoms or outcomes of SARS-CoV-2 infections; however, ACE2 levels in lungs were found to be increased in patients with comorbidities associated to severe COVID-19 clinical manifestations (24), whereas polymorphisms of ACE2 have been already described to play a role in hypertension and cardiovascular diseases (25), particularly in association with type 2 diabetes (23), all conditions predisposing to an increased risk of dying from COVID-19 (26). Despite early studies, the presence of Spike mutations potentially altering the binding with ACE2 is still largely under-investigated, as is the role of ACE2 variants in the human population in determining patient-specific molecular interactions between these two proteins.

In the present study, we aim at detecting which Spike and ACE2 AAs are the most important in determining the SARS-CoV-2 entry interaction and analyze which ones have already mutated in the population. The task is clinically relevant, providing a functional characterization of present and future mutations targeting the ACE2/Spike binding and detected by sequencing SARS-CoV-2 on a patient-specific basis. Characterizing the variability of both proteins must be taken in consideration in the process of developing anti-COVID-19 strategies, such as the Spike-based vaccine currently deployed by the National Institute of Allergy and Infectious Diseases and Moderna (27).

## Results

We set out to analyze the key AAs involved in the Spike/ACE2 interaction, in order to highlight which ones may alter the binding affinity and therefore etiological and clinical properties of different SARS-CoV-2 variants on different patients. Following that, we determined which Spike and ACE2 AA variations relevant for this interaction have been observed in the SARS-CoV-2 and human population, respectively.

### Structural analysis of Spike/ACE2 interaction

We obtained structural models of the SARS-CoV-2 Spike interacting with the human ACE2 from three recent X-ray structures, deposited on the Protein Data Bank: 6LZG (10), 6M0J (11) and 6VW1 (12). For 6VW1, two Spike/ACE2 complexes were available, so we report results for both as 6VW1-A and 6WV1-B, separately. All models show the core domains of interaction, located in the region of AA 330-530 for Spike and in the region AA 15-615 of ACE2. Full length proteins would be 1273 AAs (Spike only known isoform, from reference SARS-CoV-2 genome NC_045512.2) and 805 AAs (ACE2 isoform 1, UniProt id Q9BYF1-1).

Selected PDB entries are wild type and their primary sequence and the higher order structures were identical. Residues 517-519 were missed in 6VW1-B. With the aim to investigate the conformation variability, PDB complexes were aligned by backbone and the Root Mean Square deviation (RMSd) was computed on all equivalent not hydrogen atoms. RMSd data have shown some conformation flexibility that confirmed our idea to take into account all PDB structures in the next investigation (Fig 1).

**Figure 1.**
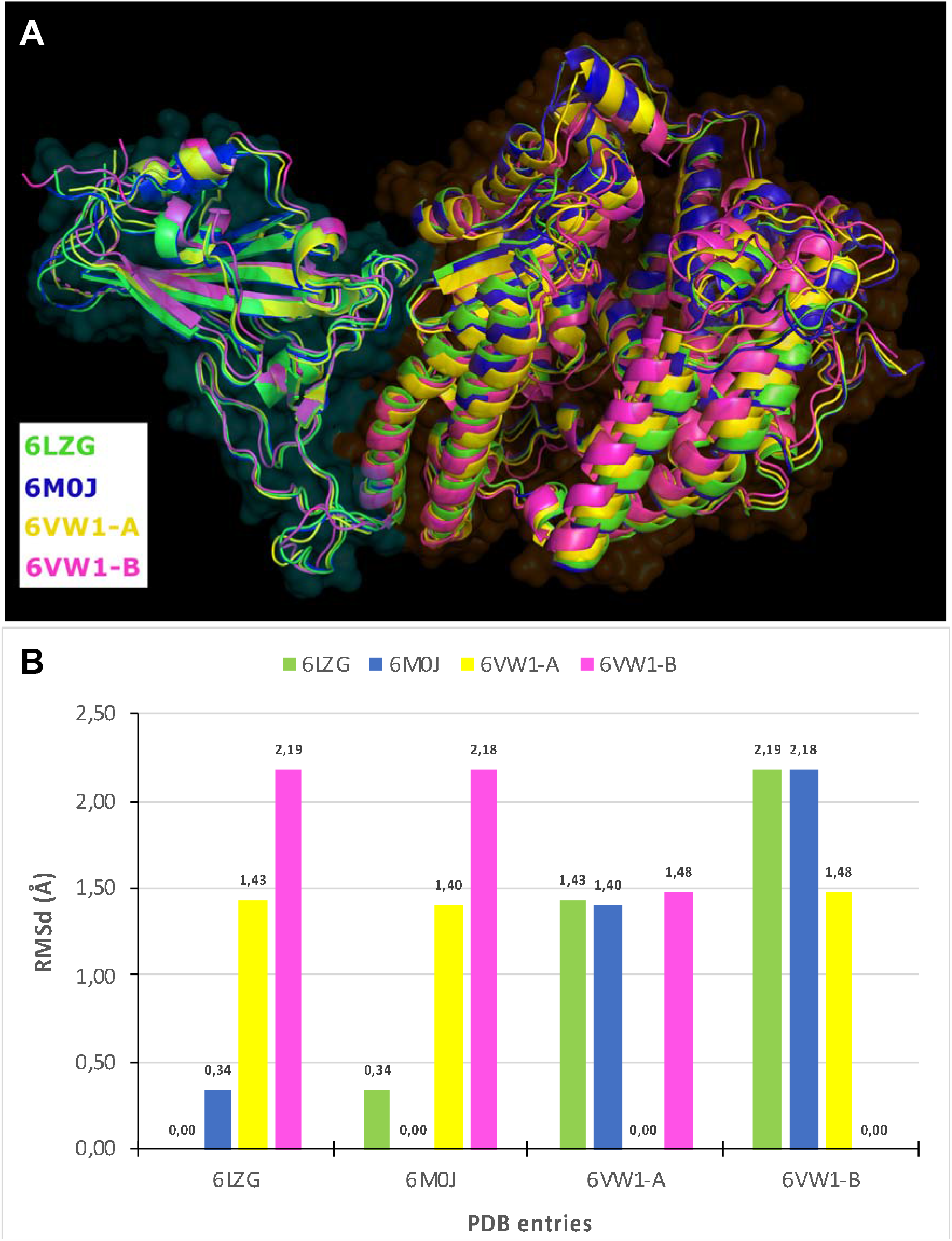
Conformational comparison of Spike-ACE2 PDB complexes: (A) alignment of PDB entries, Spike and ACE2 are respectively surrounded by cyan and orange fog, and (B) bar graph showing RMSd (in Å) computed on structures aligned without hydrogen atoms.

The GBPM method was originally developed for identifying and scoring pharmacophore and protein-protein interaction key features by combining GRID molecular interaction fields (MIFs) according to the GRAB tool algorithm (28). In the present study, GBPM has been applied to all selected complex models considering Spike and ACE2 either as host or guest. DRY, N1 and O GRID probes were considered for describing hydrophobic, hydrogen bond donor and hydrogen bond acceptor interaction. For each probe a cut-off, required for highlighting the most relevant MIFs points, was fixed above the 30% from the corresponding global minimum interaction energy value. With respect to the known GBPM application, where pharmacophore features are used for virtual screening purposes, here these data guided us in the complex stabilizing AAs identification. In fact, Spike or ACE-2 residues, within 3 Å from GBPM points, were marked as relevant in the host-guest recognition and were qualitatively scored by assigning them the corresponding GBPM energy. If a certain residue was suggested by more than one GBPM point, its score was computed as summa of the related GBPM points energy (Fig. 2).

**Figure 2.**
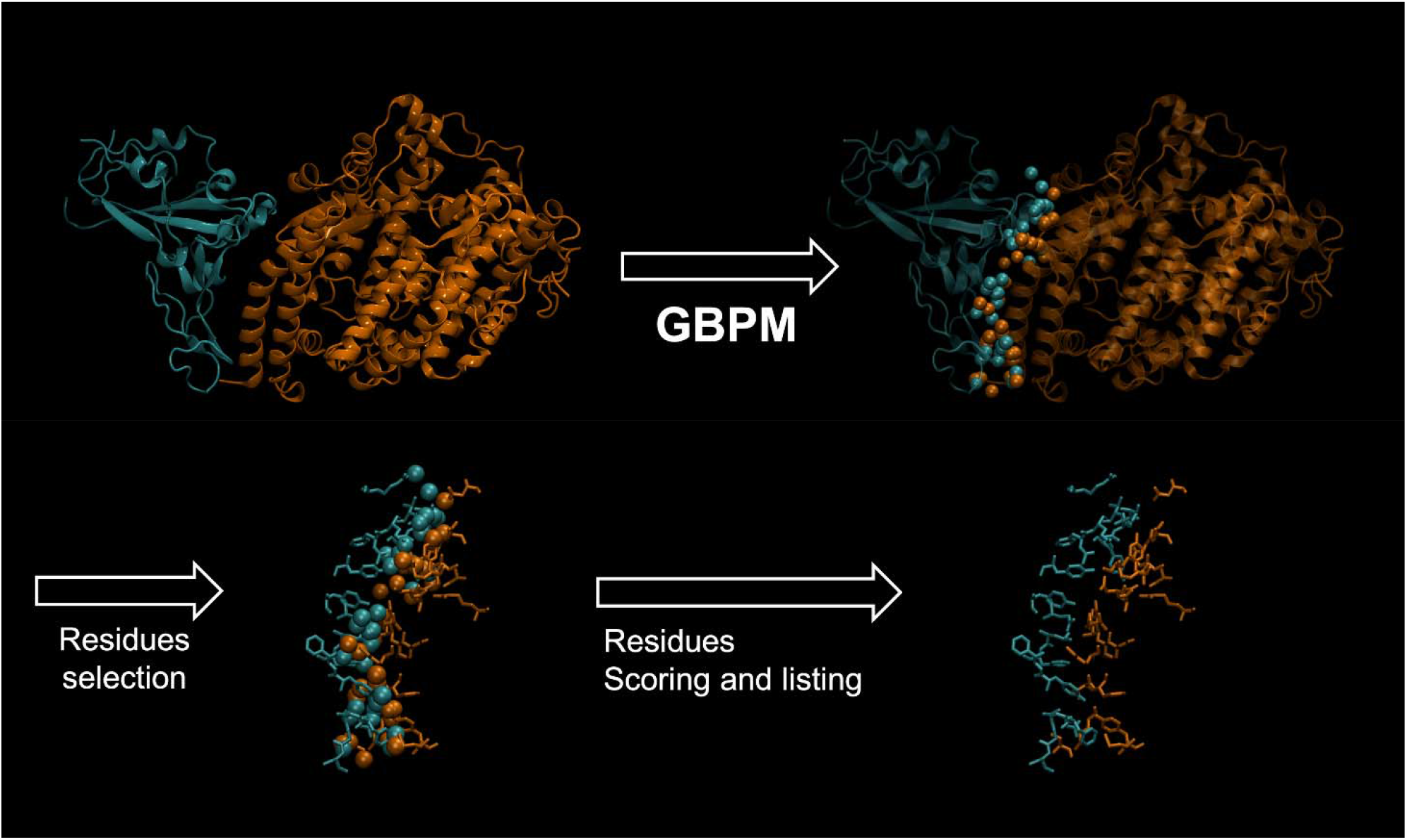
Summary of the pipeline adopted by GBPM to identify key residues contributing to the SARS-CoV-2 Spike / Human ACE2 interface. Spike is depicted in cyan, and ACE2 in orange, based on the 6LZG PDB model (10). Residues highlighted by GBPM are then tested for mutation frequency in the worldwide SARS-CoV-2 population.

Finally, for each selected residue, the four models averaged score was considered for estimating the role in complex stabilization. Taking into account their average scores, Spike and ACE2 AAs were divided by quartiles to facilitate the interpretation of the results: quartile 1 (Q1) includes the strongest complex stabilization contributors; quartile 2 (Q2) contains residues less important than those reported in Q1 but most relevant of those included in quartile 3 (Q3); quartile 4 (Q4) indicates the weakest predicted interacting AAs. Such an extension of the original approach allowed us to highlight known relevant interaction residues of both Spike (Table 1) and ACE-2 (Table 2).

**Table 1.** GBPM scores, average values, and quartile distribution of Spike relevant AAs in three PDB models. GBPM scores and average values are reported in kcal/mol.

| Residue | # | PDB entries |  |  |  | GBPM |  |
| --- | --- | --- | --- | --- | --- | --- | --- |
|  |  | 6LZG | 6M0J | 6VW1-A | 6VW1-B | Average score | Quartile |
| LYS | 417 | -43.58 | -12.12 | 0.00 | 0.00 | -13.93 | Q2 |
| ASN | 439 | 0.00 | 0.00 | -12.30 | -34.94 | -11.81 | Q2 |
| GLY | 446 | -22.52 | -5.75 | 0.00 | -10.32 | -9.65 | Q3 |
| GLY | 447 | -5.63 | 0.00 | 0.00 | 0.00 | -1.41 | Q3 |
| TYR | 449 | -25.72 | -6.38 | -20.37 | -24.76 | -19.31 | Q1 |
| TYR | 453 | 0.00 | 0.00 | -1.77 | -1.76 | -0.88 | Q4 |
| LEU | 455 | -11.59 | -16.82 | -21.78 | -7.04 | -14.31 | Q2 |
| PHE | 456 | -34.20 | -30.16 | -39.72 | -20.76 | -31.21 | Q1 |
| ALA | 475 | -52.35 | -49.72 | -38.73 | -77.00 | -54.45 | Q1 |
| GLY | 476 | -21.72 | 0.00 | -17.16 | -34.59 | -18.37 | Q2 |
| SER | 477 | -22.32 | 0.00 | -11.44 | -40.68 | -18.61 | Q2 |
| GLU | 484 | -8.52 | -13.23 | 0.00 | 0.00 | -5.44 | Q3 |
| PHE | 486 | -28.99 | -53.63 | -32.56 | -53.43 | -42.15 | Q1 |
| ASN | 487 | -31.67 | -59.57 | -33.98 | -52.21 | -44.36 | Q1 |
| TYR | 489 | -62.10 | -27.67 | -45.92 | -69.38 | -51.27 | Q1 |
| PHE | 490 | -4.58 | -4.48 | -22.90 | -40.32 | -18.07 | Q2 |
| GLN | 493 | -37.20 | -56.08 | -79.60 | -70.51 | -60.85 | Q1 |
| GLY | 496 | -15.54 | -8.74 | -18.72 | -16.80 | -14.95 | Q2 |
| PHE | 497 | -8.86 | 0.00 | -4.68 | -29.10 | -10.66 | Q3 |
| GLN | 498 | -77.24 | -80.38 | -42.34 | 0.00 | -49.99 | Q1 |
| PRO | 499 | 0.00 | 0.00 | 0.00 | -11.64 | -2.91 | Q3 |
| THR | 500 | 0.00 | -66.00 | -92.90 | -122.50 | -70.35 | Q1 |
| ASN | 501 | -60.14 | -61.04 | -61.82 | -70.59 | -63.40 | Q1 |
| GLY | 502 | -24.84 | -35.42 | -39.45 | -40.92 | -35.16 | Q1 |
| VAL | 503 | 0.00 | -5.37 | -5.45 | -5.54 | -4.09 | Q3 |
| TYR | 505 | -30.60 | -23.22 | -20.90 | -40.62 | -28.84 | Q1 |

**Table 2.** GBPM scores, average values, and quartile distribution of ACE2 relevant AAs in three PDB models. GBPM scores and average values are reported in kcal/mol.

| Residue | # | PDB entries |  |  |  | GBPM |  |
| --- | --- | --- | --- | --- | --- | --- | --- |
|  |  | 6LZG | 6M0J | 6VW1-A | 6VW1-B | Average Score | Quartile |
| SER | 19 | -31.45 | -26.08 | -53.61 | -79.33 | -47.62 | Q1 |
| GLN | 24 | -31.15 | -23.62 | -34.15 | -85.23 | -43.54 | Q1 |
| THR | 27 | -16.93 | -32.58 | -38.70 | -16.65 | -26.22 | Q2 |
| PHE | 28 | -20.68 | -25.02 | -14.10 | -27.48 | -21.82 | Q2 |
| ASP | 30 | 0.00 | -17.01 | 0.00 | 0.00 | -4.25 | Q3 |
| LYS | 31 | -84.06 | -43.67 | -32.98 | -46.60 | -51.83 | Q1 |
| HIS | 34 | 0.00 | -30.42 | -27.78 | -67.56 | -31.44 | Q2 |
| GLU | 35 | -11.73 | 0.00 | 0.00 | -19.40 | -7.78 | Q2 |
| GLU | 37 | -11.58 | -20.36 | -11.83 | -20.52 | -16.07 | Q2 |
| ASP | 38 | -41.09 | -40.52 | -25.75 | -34.16 | -35.38 | Q2 |
| TYR | 41 | -52.50 | -75.07 | -62.35 | -76.07 | -66.50 | Q1 |
| GLN | 42 | -36.78 | -37.15 | -28.53 | -63.49 | -41.49 | Q2 |
| LEU | 45 | -12.80 | -16.43 | 0.00 | -16.20 | -11.36 | Q2 |
| LEU | 79 | 0.00 | 0.00 | 0.00 | -5.99 | -1.50 | Q3 |
| MET | 82 | 0.00 | 0.00 | -6.36 | -6.00 | -3.09 | Q3 |
| TYR | 83 | -40.50 | -66.29 | -57.86 | -60.81 | -56.37 | Q1 |
| GLU | 329 | 0.00 | 0.00 | 0.00 | -17.25 | -4.31 | Q3 |
| ASN | 330 | -11.84 | -5.92 | -11.82 | -6.04 | -8.91 | Q2 |
| GLY | 352 | -1.97 | -8.36 | -8.86 | -14.66 | -8.46 | Q2 |
| LYS | 353 | -79.38 | -70.11 | -120.73 | -46.03 | -79.06 | Q1 |
| GLY | 354 | -21.87 | -31.15 | -12.74 | -15.25 | -20.25 | Q2 |
| ASP | 355 | -68.95 | -81.24 | -57.99 | -89.12 | -74.33 | Q1 |
| ARG | 357 | 0.00 | -4.99 | 0.00 | 0.00 | -1.25 | Q3 |
| ALA | 386 | 0.00 | 0.00 | -4.85 | 0.00 | -1.21 | Q4 |
| ARG | 393 | 0.00 | 0.00 | -4.85 | 0.00 | -1.21 | Q4 |

Basically, the same number of AAs was highlighted for Spike (26 AAs) and ACE2 (25 AAs). The average score was also in the same range. Spike reported a population of Q1 larger than ACE2: 12 and 7 AAs, respectively. The opposite scenario was observed in the Q2 that accounted for 7 residues for Spike and 11 for ACE2. No remarkable difference can be addressed to the Q3 and Q4 Spike-ACE2 comparison. We reasoned that mutations and variants in Q1 residues could have a more relevant impact in the complex stability.

The analysis of all designed GBPM suggested the Spike - ACE2 molecular recognition is largely sustained by polar interactions, such as hydrogen bonds, and by very few putative hydrophobic contributions (Table 3).

**Table 3.**
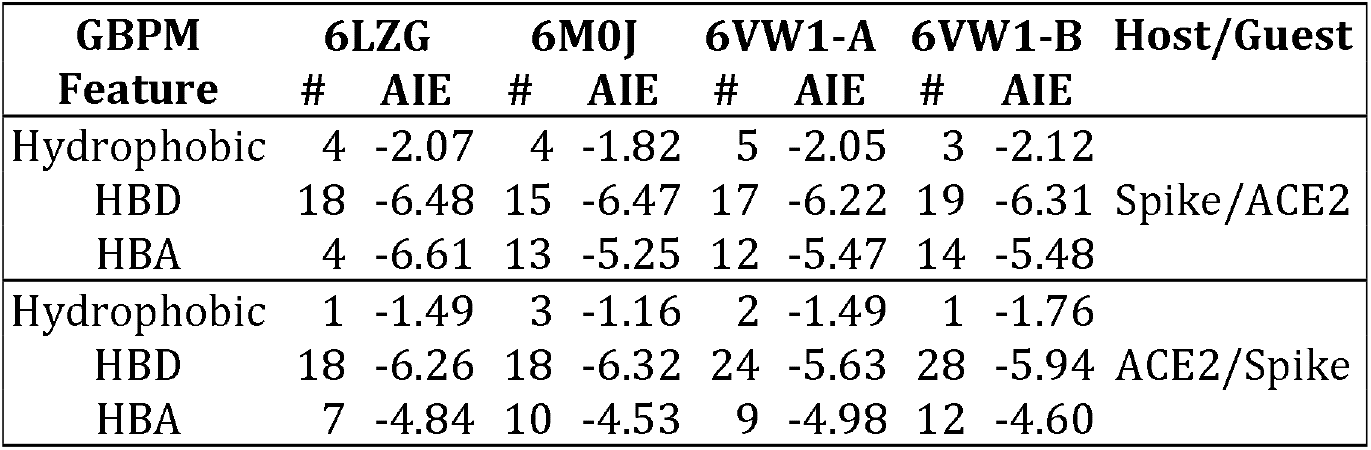
Composition of the GBPM models designed. HBD = Hydrogen Bond Donor; HBA = Hysdrogen Bond Acceptor; # = number of features; AIE = Average Interaction Energy (in kcal/mol).

| GBPM<br>Feature | 6LZG |  | 6M0J |  | 6VW1-A |  | 6VW1-B |  | Host/Guest |
| --- | --- | --- | --- | --- | --- | --- | --- | --- | --- |
|  | # | AIE | # | AIE | # | AIE | # | AIE |  |
| Hydrophobic | 4 | -2.07 | 4 | -1.82 | 5 | -2.05 | 3 | -2.12 | Spike/ACE2 |
| HBD | 18 | -6.48 | 15 | -6.47 | 17 | -6.22 | 19 | -6.31 |  |
| HBA | 4 | -6.61 | 13 | -5.25 | 12 | -5.47 | 14 | -5.48 |  |
| Hydrophobic | 1 | -1.49 | 3 | -1.16 | 2 | -1.49 | 1 | -1.76 | ACE2/Spike |
| HBD | 18 | -6.26 | 18 | -6.32 | 24 | -5.63 | 28 | -5.94 |  |
| HBA | 7 | -4.84 | 10 | -4.53 | 9 | -4.98 | 12 | -4.60 |  |

### Mutational analysis of SARS-CoV-2 Spike

We analyzed 295,507 publicly available SARS-CoV-2 full-length genome sequences collected worldwide and deposited on the GISAID database on December 30, 2020 (29). From these, we obtained 257,434 samples containing at least one AA-changing mutation in the Spike protein. A total of 3,314 different AA-changing mutations were detected in the 1,279 AA-long Spike sequence. However, many of these are unique events (or possibly even sequencing errors), as only 2,023 mutations were found in more than one sample, 788 were found in more than ten samples, and 196 in more than one hundred samples (Supplementary File 1).

We then focused on mutations located in the Spike RBD (aa 330-530) with predicted interaction contribution, as assessed by our GBPM method. The majority of mutations here are found in only a handful of samples (Table 4 and Fig 4 A), with a few notable exceptions. The mutations S477N and N439K are the most frequent in the current population and were identified in 16,547 patients (5.60%) and 5,587 patients (1.89%) respectively. These two variants (N439K and S477N) are also amongst the top 20 most frequent in the population and involve two positions productively contributing to the interaction between Spike and ACE2, according to GBPM (see Table 1 and Fig 3 for locations 439 and 477).

**Table 4.** Spike mutations located within the RBD (AA 330-530) with at least two cases in the population and non-zero GBPM average score in the ACE2/Spike interaction models. The asterisk (*) indicates a stop codon. A lower GBPM score indicates a stronger effect in the ACE2/Spike interaction.

| Mutation | Position | Abundance | Frequency | GBPM |  |
| --- | --- | --- | --- | --- | --- |
|  |  |  |  | Average Score | Quartile |
| S477N | 477 | 16547 | 0.055995 | -18.61 | Q2 |
| N439K | 439 | 5587 | 0.018906 | -11.81 | Q2 |
| N501Y | 501 | 4921 | 0.016653 | -63.3975 | Q1 |
| Y453F | 453 | 917 | 0.003103 | -0.8825 | Q4 |
| E484K | 484 | 352 | 0.001191 | -5.4375 | Q3 |
| K417N | 417 | 260 | 0.00088 | -13.925 | Q2 |
| S477I | 477 | 157 | 0.000531 | -18.61 | Q2 |
| G446V | 446 | 58 | 0.000196 | -9.6475 | Q3 |
| F490S | 490 | 53 | 0.000179 | -18.07 | Q2 |
| S477R | 477 | 49 | 0.000166 | -18.61 | Q2 |
| N501T | 501 | 47 | 0.000159 | -63.3975 | Q1 |
| L455F | 455 | 44 | 0.000149 | -14.3075 | Q2 |
| G476S | 476 | 43 | 0.000146 | -18.3675 | Q2 |
| E484Q | 484 | 43 | 0.000146 | -5.4375 | Q3 |
| A475V | 475 | 35 | 0.000118 | -54.45 | Q1 |
| F486L | 486 | 34 | 0.000115 | -42.1525 | Q1 |
| F490L | 490 | 18 | 6.09E-05 | -18.07 | Q2 |
| YQ505WK | 505 | 14 | 4.74E-05 | -28.835 | Q1 |
| Q493L | 493 | 12 | 4.06E-05 | -60.8475 | Q1 |
| V503F | 503 | 9 | 3.05E-05 | -4.09 | Q3 |
| E484A | 484 | 8 | 2.71E-05 | -5.4375 | Q3 |
| G446S | 446 | 7 | 2.37E-05 | -9.6475 | Q3 |
| E484D | 484 | 4 | 1.35E-05 | -5.4375 | Q3 |
| Q493* | 493 | 4 | 1.35E-05 | -60.8475 | Q1 |
| Y505W | 505 | 4 | 1.35E-05 | -28.835 | Q1 |
| G476A | 476 | 3 | 1.02E-05 | -18.3675 | Q2 |
| S477G | 477 | 3 | 1.02E-05 | -18.61 | Q2 |
| F456L | 456 | 2 | 6.77E-06 | -31.21 | Q1 |
| V503I | 503 | 2 | 6.77E-06 | -4.09 | Q3 |
| Y449F | 449 | 2 | 6.77E-06 | -19.3075 | Q1 |

**Figure 3.**
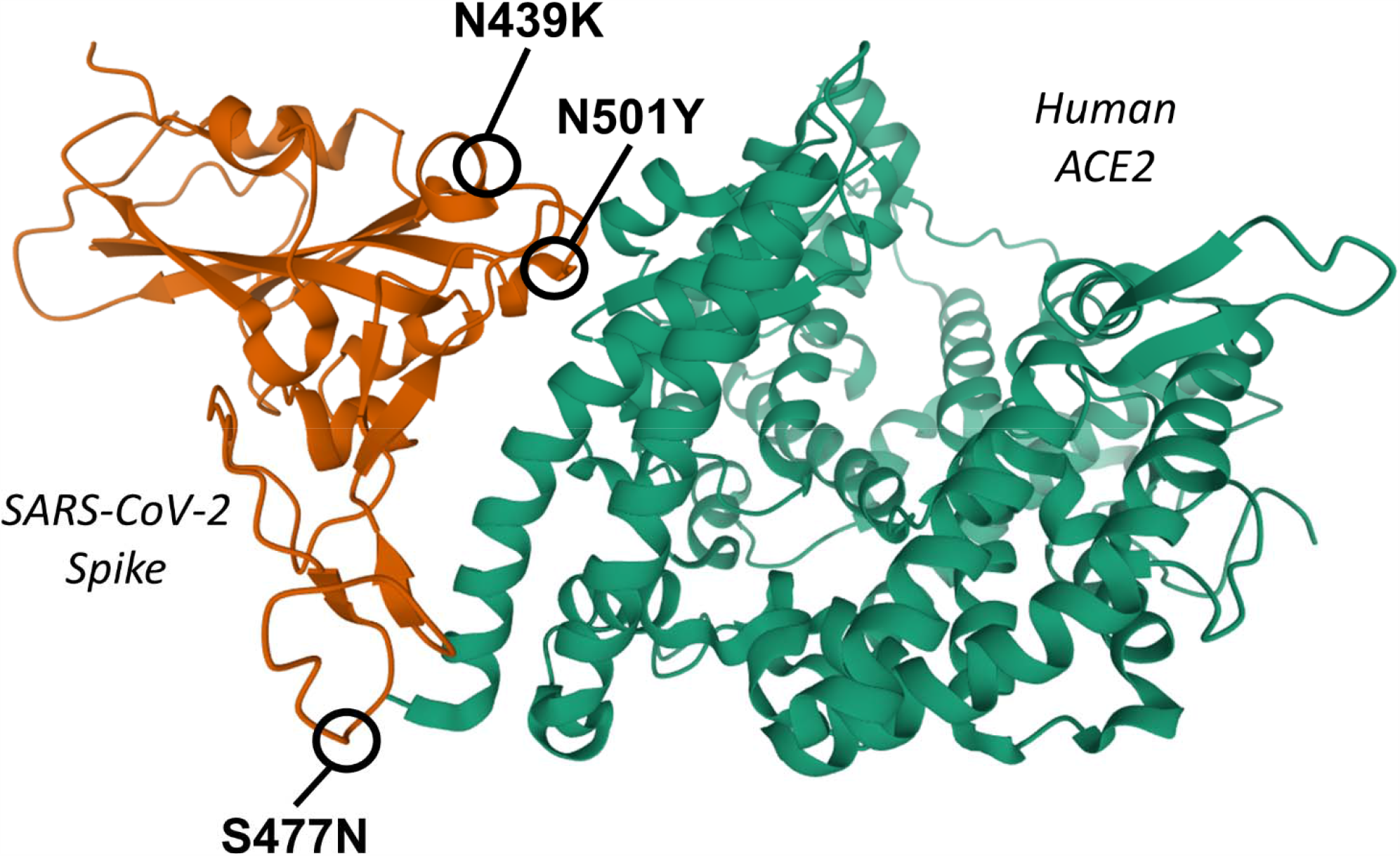
3D ribbon representation of the interaction domains of SARS-CoV-2 Spike (left, orange) and human ACE2 (right, green), based on the crystal structure 6LZG deposited on Protein Data Bank and produced by (10). The positions of the three most frequent Spike mutations in the interacting region (AA 350-550) with a non-zero GBPM score are indicated: N439K, N501Y and S477N.

The graphical inspection of the PDB structures revealed that Spike Asparagine (N) 439, raked at GBPM Q2, is mainly involved in intra-protein interaction. In fact, by means of its backbone sp2 oxygen atom, N439 accepts one hydrogen bond from Spike Serine 443 sidechain and, by its sidechain amide group, donates one hydrogen bond to the Spike Proline 499 backbone: all these AAs are located into a random coil loop of Spike so the N439K could minimally modify the Spike-ACE2 recognition. On the other hand, after the theoretical mutation of the Asparagine 439 with a Lysine, it is possible to predict a productive electrostatic interaction between the new net positively charged residue and the ACE2 Glutamate 329. Such a long-distance interaction could improve the stabilization of the complex with respect to the Spike wild type (Figure S1).

A similar effect could be addressed to the mutation at position 477. Serine (S) 477 is a weak contributor to the complex interaction. In all PDB entries we selected, Serine 477 is located into a solvent exposed random coil loop. No interaction with ACE2 or Spike residues can be observed. Actually, the GBPM analysis included such a residue in Q2. Conversely, its mutation to Asparagine (S477N), in our *in silico* model, revealed the possibility to establish hydrogen bond to the ACE2 Serine 19 that can clearly result in a stabilization of the complex (Figure S2). Moreover, position 477 is also affected by three other events with lower occurrence: S477I, S477R and S477G, with 6, 2 and 2 observations (Table 4). Among all, the S447R could be the most interesting one. Actually, a net positively charged residue, such as Arginine (R), can establish a weak electrostatic interaction to ACE2 Glutamate 87, as suggested by a theoretical model we built. The S477I and S477G could modify the conformation of a random coil segment, so it does not appear very relevant. Conversely, S477N and S477G could productively contribute to the Spike ACE2 complex stabilization. Of course, deeper theoretical and experimental investigations should be carried out to confirm this hypothesis. Unfortunately, full-scale simulations cannot be rigorously performed today because the available 3D structural models report only fragments of the complex between Spike and ACE2.

The third most common mutation, N501Y (Fig 3), targets an AA predicted to have a strong role in the interaction in all four models, sitting in the GBPM Q1. N501Y was detected in 4,921 patients (1.67% of the dataset): the majority of which were located in the United Kingdom (29). From a structural point of view, we predict that a substitution, at position 501, of an Asparagine (N) with a Tyrosine (Y) may have an effect: their Total Polar Surface Area (TPSA), equal to 101.29 and to 78.43 Å 2 respectively, is different, however both their sidechains can donate/accept a hydrogen bond. Therefore, their contribution to complex stabilization may be slightly different, also taking into account the chemical environment. In fact, the wild type Asparagine 501 donates one hydrogen bond to ACE2 Tyrosine 41: such an interaction could be possible also for N501Y mutant or, as we observed in our theoretical model, it could be replaced by pi-pi stacking (Figure S3). The rapid increase in frequency of mutation N501Y has been recently observed in the United Kingdom and other countries, as it is one of the variants characterizing lineage B1.1.7 (30). The Asparagine/Tyrosine substitution in Spike position 501 could contribute to determine an evolutionary advantage for this lineage, based on differential affinity for the human receptor ACE2 (31, 32).

A less frequent mutation amongst those predicted to contribute to the ACE2/Spike interaction is G476S, detected in 43 samples (0.02%), and supported by three out of four structural models (Table 1, Fig 4 B). The Glycine (G) 476 was included by GBPM analysis in Q2: its contribution to the complex stabilization is weak. Conversely to the other mutation described here, the replacement of Glycine 476 with a Serine (S) could have more evident effects on Spike ACE2 molecular recognition. In fact, in all PDB entries, the alpha carbon of this Glycine is very close, about 4 Å, to the sidechain amide group of the ACE2 Glutamine 24. Between these two AAs no productive interaction can be established but the substitution of the Spike Glycine with a Serine could allow one inter-protein hydrogen bond to ACE2 Glutamine 24. Moreover, G476S could establish the same interaction with Spike Glutamine 478 that could stabilize the conformation of a random coil segment of the viral protein resulting in a better pre-organization to the ACE2 recognition (Figure S4).

**Figure 4.**
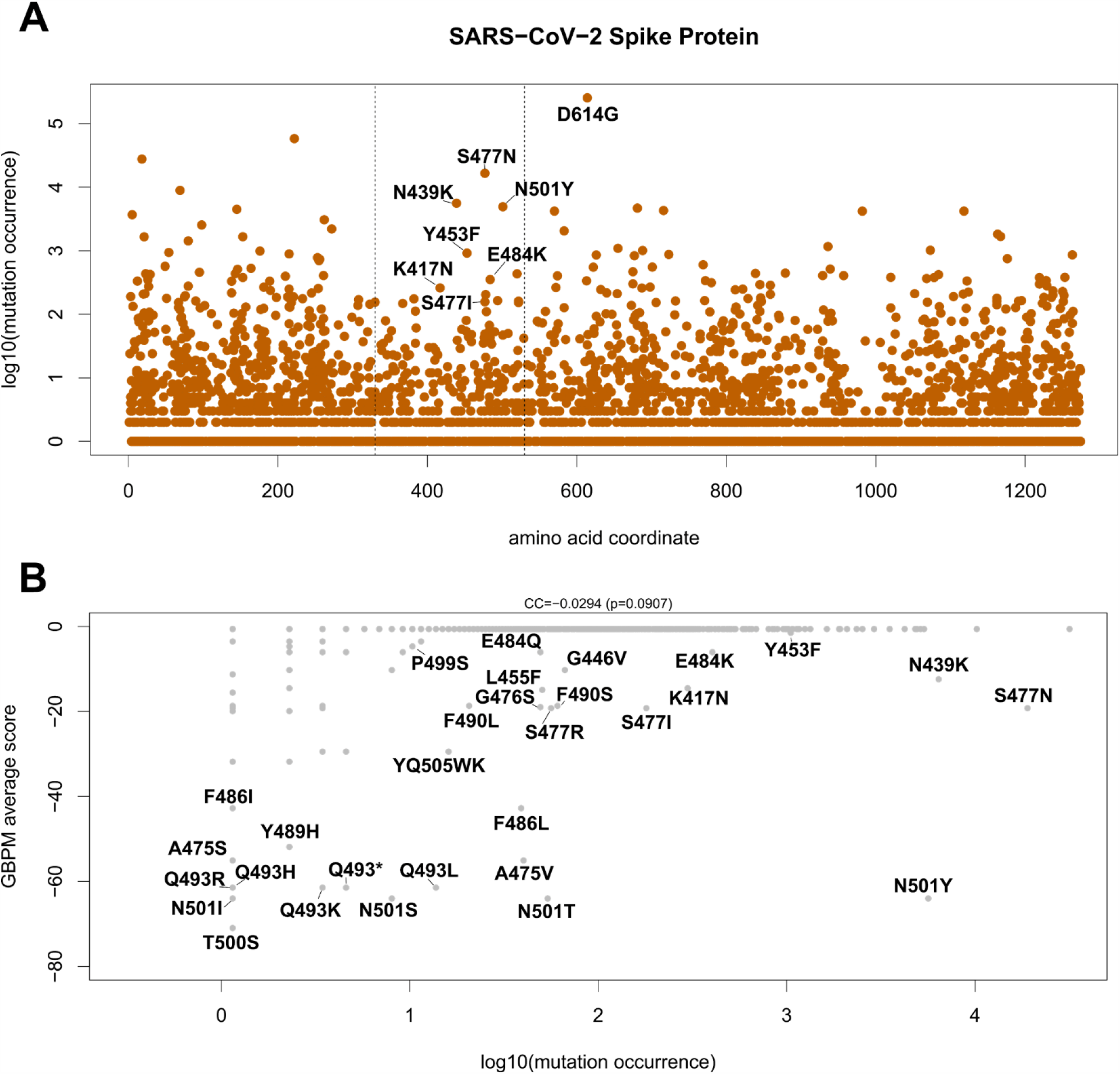
(A) Occurrence of AA-changing variants on SARS-CoV-2 Spike protein. X-axis indicates the position of the affected AA. Y-axis indicates the log10 of the number of occurrences of the variant in the SARS-CoV-2 dataset. Labels indicate variants affecting ACE2/Spike binding and detected in at least 5 SARS-CoV-2 sequences. Vertical dashed lines indicate crystalized region analyzed (aa 330 – 530). The D614G variant, located outside the RBD, is also indicated. (B) Scatter plot indicating the occurrence of the variant in the population (x-axis) and the GBPM score of the reference AA in the model (y-axis). Mutations with non-zero GBPM score are indicated. CC indicates the Pearson correlation coefficient and p indicates the p-value of the CC.

Another Spike residue, predicted by our analysis for playing a relevant role in ACE2 recognition, is the Glutamine 493 (Table 1). The GISAID data revealed that such an aminoacid is rarely replaced by a Leucine (Q493L) or by an Arginine (Q493R). These mutations could affect the recognition of ACE2 in an opposite way. Spike Glutamine 493 is involved in hydrogen bond with ACE2 Glutamate 35. The mutation Q493L cannot establish such a productive contribution and could only hydrophobically interact to Spike Leucine 455. Conversely, Q493R could locate its net positively charged sidechain into an ACE2 pocket delimited by Aspartate 30, Histidine 34 and Glutamate 35. Such a positioning could produce a remarkable electrostatic stabilization of the complex (Figure S5).

In general, we could observe that AAs with the strongest evidence for interaction contribution in the Spike/ACE2 interface tend not to diverge from the reference (Fig 4 B), which may indicate a solid evolutionary constraint to maintain the interface residues unchanged. For example, one of the most relevant 1^st^ quartile AA in the ACE2/Spike interaction, Glutamine (Q) 493, is rarely mutated, with 12 cases of Q493L, 4 of Q493* (the substitution of Q493 with a stop codon), 3 of Q493K, and 1 of Q493R and Q493H. One possible exception is the aforementioned Spike mutation N501Y, located in the strongest 1^st^ quartile GBPM-predicted AA for ACE2 binding, which was found in the considerable number of 4921 different patients.

### Mutational analysis of human ACE2

We also investigated the variants of human ACE2, since these could constitute the basis for patient-specific COVID-19 susceptibility and severity. ACE2 protein sequence is highly conserved across vertebrates (33) and also within the human species (34), with the most frequent missense mutation (rs41303171, N720D) present in 1.5% of the world population (Supplementary File 2).

Our analysis shows that only 5 variants of ACE2 detected in the human population are also located in the ACE2/Spike direct binding interface (Table 5 and Fig 5). Of these, rs73635825 (causing a S19P AA variant) is both the most frequent in the population (0.06%) and the most relevant in the interaction with the viral protein, with a GBPM score of−47.6175 (Q1) and support from all 4 models (Table 2). The rs73635825 SNP frequency is higher in the population of African descent (0.2%). The second SNP, rs143936283 (E329G, Table 5) is a very rare allele (0.0066%) in the European (non-Finnish) Asian population. The rs766996587 (M82I) SNP is also a very rare allele (0.0066%) found in the African population. E37K (rs146676783) is more frequent in the Finnish (0.03%) and G352V (rs370610075) in the European non-Finnish (0. 007%) population. None of these five SNPs have a reported clinical significance, according to dbSNP and literature search (35).

**Table 5.** ACE2 variants with non-zero GBPM score in the Spike interaction model.

| variant | rsID | Allele Frequency | GBPM |  |
| --- | --- | --- | --- | --- |
|  |  |  | Average Score | Quartile |
| S19P | rs73635825 | 0.000655 | -47.62 | Q1 |
| E329G | rs143936283 | 6.63E-05 | -4.31 | Q3 |
| M82I | rs766996587 | 6.62E-05 | -3.09 | Q3 |
| E37K | rs146676783 | 5.68E-05 | -16.07 | Q2 |
| G352V | rs370610075 | 3.8E-05 | -8.46 | Q2 |

**Figure 5.**
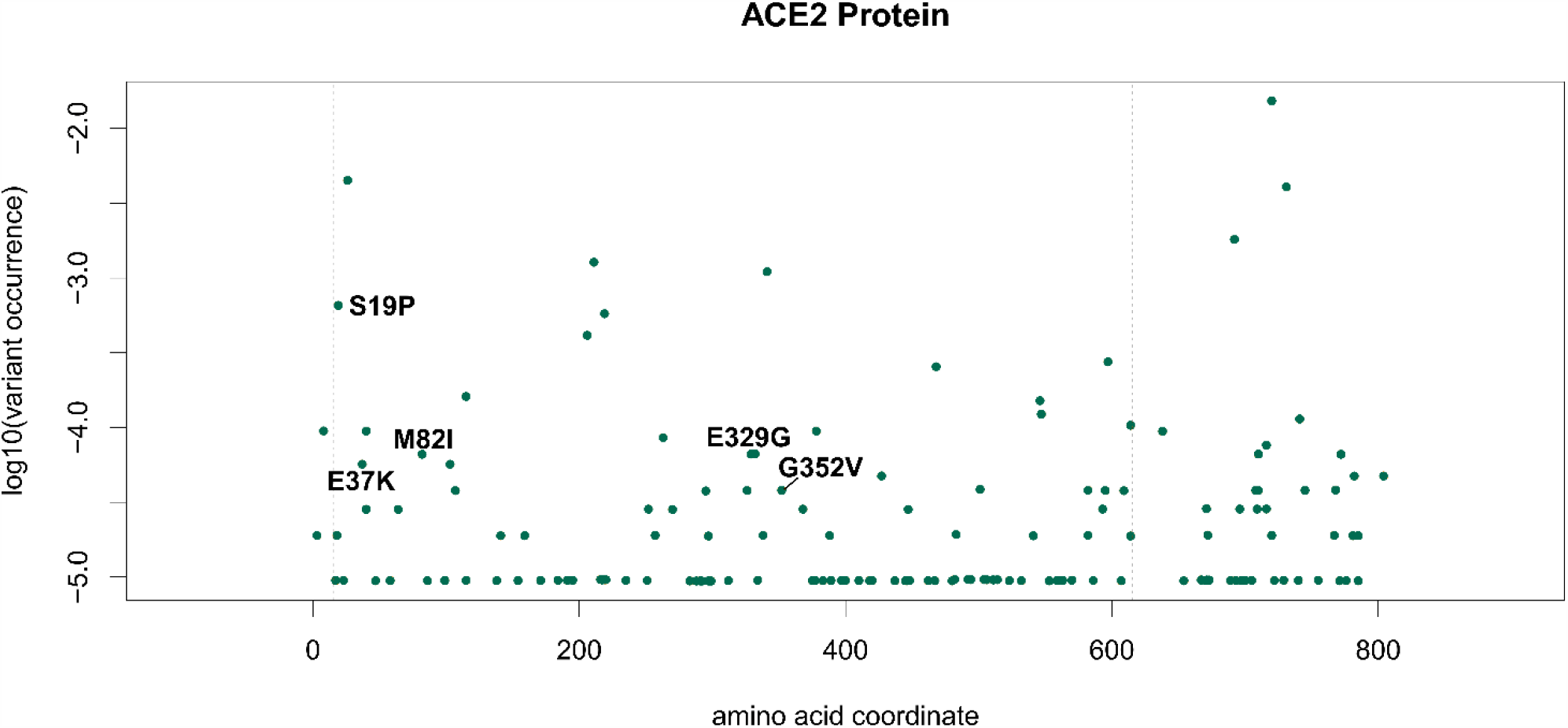
Frequency of mutations on ACE2. X-axis indicates the AA position in isoform 1 (UniProt id Q9BYF1-1). Y-axis indicates the allele frequency in the global population according to the GNOMAD v3 database. Labels indicate AA changes observed in the human population with non-zero GBPM average score in the ACE2/Spike interaction models. Vertical dashed lines indicate the crystalized region analyzed in this study (aa 15 – 615).

It must be mentioned that M82I, together with S19P, has been predicted to adversely affect ACE2 stability (36). M82I, together with E329G, has been simulated to increase binding affinity with Spike when compared to wild type ACE2, hypothesizing greater susceptibility to SARS-CoV-2 for patients carrying these variants (37). Instead, E37K (37) and G352V (38) were predicted to possess a lower affinity with Spike, suggesting lower susceptibility to the infection. However, while describing potential explanations to the existence of a possible predisposing genetic background to infection, all these studies remain inconclusive in linking allele variants to COVID-19 susceptibility.

Structurally, the S19P variant may greatly differ from the reference sequence in the interaction with ACE2: Serine (S) is a polar residue, able to accept and donate, by means of its side chain alcoholic group, a hydrogen bond. Proline (P), on the other hand, cannot be involved in hydrogen bonding, and therefore should establish a weaker interaction with Spike. In fact, ACE2 Serine 19 sidechain donates a hydrogen bond to Spike Alanine 475 backbone (Figure S6) and potentially could establish the same interaction with Spike Glycine (G) 476, which could also be mutated (Table 4). Both Methionine (M) 82 and Glutamate (E) 329 are in Q3 minimally contributing to Spike ACE2 recognition (Figures S7 and S8). They are located within two alpha helices so their mutation could modify the secondary structure of ACE2 corresponding to a different affinity against Spike. Such a possibility should be more evident in the case of E329G because Glutamate 329 sidechain is involved in hydrogen bond with ACE-2 Glutamine 325.

## Discussion

SARS-CoV-2 Spike evolved through a series of adaptive mutations that increased its affinity for the human ACE2 receptor (39). There is no reason to believe that the evolution and adaptation of the virus will stop, making continuous sequencing and mutational tracking studies of paramount importance to strategically contain COVID-19 (40). In our study, we highlighted which specific locations of Spike can influence the ACE2 molecular recognition, required for the viral entry into the host cell (5). We further showed that some mutations are already present in the SARS-CoV-2 population that may weakly affect the interaction with the human receptor, specifically Spike N439K, S477N and N501Y. These mutations are rising in the viral population (>1%) and in particular N501Y is one of the key mutations characterizing lineage B.1.1.7 (32), which has seen a recent dramatic increase in frequency in the United Kingdom (30). Having identified this mutation proves that our combination of targeted mutation frequency and GBPM is a useful pipeline to monitor events in the key region used by SARS-CoV-2 to recognize and enter human bronchial cells. The same approach can be used to monitor, in the future, if any of these events will increase in frequency, suggesting an adaptation to the human host leveraging a higher affinity with ACE2.

On the other hand, we studied the variants in the human ACE2 population, identifying 5 loci that can affect the binding with SARS-CoV-2 Spike. They are all rare variants, with the most frequent, S19P, present in 0.06% of the population, and with no known clinical significance. However, other *in silico* studies have predicted their role in decreasing ACE2 stability (S19P and M82I) (36), and in altering the affinity with Spike (increasing it: M82I and E329G (37); decreasing it: E37K (37) and G352V (38)). The most common ACE2 variant, rs41303171 (N720D), is not located in the binding region, and so far its predicted effects on the etiopathology of COVID-19 are still largely conjectural and associated to neurological complications via mechanisms probably independent from direct interaction with Spike (41).

It remains to be seen whether, in the future, the combination of Spike and ACE2 sequences will produce novel and unexpected COVID-19 specificities, that will require granular efforts in developing wider-spectrum anti-SARS-CoV-2 strategies, such as vaccines or antiviral drugs. So far, our analysis has shown a location on the Spike/ACE2 complex where both proteins vary in the viral/human population, specifically on ACE2 S19 and Spike A475/G476. While, as described in our Results, these mutations on Spike are not likely to strongly affect the interaction surface, future combinations of ACE2/Spike variants may have peculiar effects that will require constant mutation monitoring. Identifying single or multiple AAs involved in this viral entry interaction will allow for personalized diagnosis and clinical prediction based on the specific combination of SARS-CoV-2 strain and ACE2 variant. Personalized COVID-19 treatment will require targeted sequencing of the patient ACE2 and Spike, to identify the combination causing the specific case. This technical obstacle can be further complicated by the intra-host genetic variability of SARS-CoV-2, which has recently been reported from RNA-Sequencing studies (42).

Structural investigation will benefit, in the next future, from the availability of experimental structural models reporting the complete sequence of both Spike and ACE2, or at least Spike. This will allow more rigorous computational analyses (i.e. molecular dynamics simulation, free energy perturbation) on the effect of mutations on the Spike/ACE2 recognition. Beyond the complex investigated in this manuscript, our approach can be fully extended to any other partners in the SARS-CoV-2/human interactome, for example the recently discovered interaction between viral protease NSP5 (43) and human histone deacetylase HDAC2 (44), which is indirectly responsible for the transcriptional activation of pro-inflammatory genes. Our approach can also be extended to other viruses exploiting human receptors as an entry mechanism, such as CD4 for the Human Immunodeficiency Virus (HIV) or TIM-1 for the Ebola virus (45).

## Materials and Methods

### Structural analysis

The PDB (46) was searched for high resolution Spike/ACE2 complexes. PDB entries 6LZG (10), 6M0J (11) and 6VW1 (12), reporting the Spike RBD interacting to ACE2, have been retrieved and taken into account for our GBPM analysis (28). Such a computational approach compares GRID (47) molecular interaction fields (MIFs) computed on a generic complex (A) and on its host (B) and guest (C) components, separately. Actually, MIFs describe the interaction between a certain probe and a certain target. If the target is represented by a complex, depending on the selected area, the MIF energies can be referred to the interaction between the probe and one of the complex subunits or, at the host/guest interface, with both of them. The GBPM analysis, objectively, highlights these last. Five steps are required: (1) the complex A is disassembled in its subunits B and C; (2) MIFs are computed on A, B and C by using the most appropriate GRID probes. A hydrogen bond acceptor/donor and a generic hydrophobic probe can describe the basic interaction. Because GRID MIFs are stored as a 3D matrix of interaction energy points (IEP), the same box dimensions are adopted in all calculations; (3) each IEP of B is compared with respect to the equivalent point of A generating a new MIFs named D. The following algorithm, available into the GRAB tool, is applied: if IEP(A) > 0 and IEP(B) > 0 then IEP(D) = 0; if IEP(A) > 0 and IEP(B) < 0 then IEP(D) = IEP(B); if IEP(A) < 0 and IEP(B) > 0 then IEP(D) =−IEP(A); if IEP(A) < 0 and IEP(B) < 0 then IEP(D) = IEP(A)-IEP(B). The resulting MIF D reports as negative energy values the productive interaction between the GRID probe and B and the interface A and B; (4) in order to obscure the interaction between the probe and B, MIFs D and C are compared, by using the GRAB approach, producing to a new MIF E; (5) the most relevant interaction points (GBPM features) of the MIF E are, finally, selected taking into account an energy cutoff 15% above the global minimum. Supplementary figures focusing on the most relevant mutation are available in Supplementary File 3.

Before starting the GBPM analysis, co-crystalized water molecules were removed from PDB structures. In 6VW1, showing two Spike-ACE2 complexes, namely chains A-E and B-F, both structures have been investigated and further reported as model A and B, respectively. All selected complexes have been conformationally compared one each other by alignment and computing the RMSd on the cartesian coordinates of equivalent not hydrogen atoms. DRY, N1 and O original GRID probes have been used to highlight hydrophobic, hydrogen bond donors and acceptors areas. In order to identify the most relevant residues of both Spike and ACE2, we conceptually and technically extended the GBPM algorithm, originally designed for drug/target interactions (28). In the GBPM analysis presented here, the two interacting proteins have been considered either as host and guest units, and relevant AAs were selected if their distance from GBPM features was lower or equal to 3 Å. For each PDB model, the selected residues were scored as summa of the corresponding GBPM features interaction energy.

In order to prevent unrealistic distortion of the Spike-ACE2 complex, due the usage of structures not covering the full length of the interacting proteins, the mutations effect has been qualitatively estimated by means of the mutagenesis tool implemented in PyMol software (48). Wild type residues have been replaced by the mutation and the new sidechain conformations have been optimized taking into account the neighboring AAs. The graphical analysis was carried out onto the predicted most populated rotamers. On the basis of its better X-ray resolution, the 6M0J PDB structure has been selected for the above reported investigation.

### Genetical analysis

SARS-CoV-2 genome sequences from human hosts and accounting for a total of 145,201 submissions were obtained from the GISAID database on 15 October 2020 (29). Low quality (with more than 5% uncharacterized nucleotides) and incomplete (<29,000 nucleotides, based on a total reference length of 29,903) sequences were removed. The resulting 135,591 genome sequences were aligned on the reference SARS-CoV-2 Wuhan genome (NCBI entry NC_045512.2) using the NUCMER algorithm (49). Position-specific nucleotide differences were merged for neighboring events and converted into protein mutations using the *coronapp* annotator (17). The results were further filtered for AA-changing mutations targeting the Spike protein.

ACE2 variants in the human population were extracted from the gnomAD database, v3, 18 July 2020 (50). We considered only missense variants affecting specific AAs in the protein sequence, for a total of 155 entries (Supplementary File 2). Graph generation was performed with the R statistical software and the *corto* package v1.1.2 (51).

## Acknowledgments

We thank the Italian Ministry of Education and Research for their financial support under the Montalcini initiative. We thank Prof. Giovanni Perini for his continued support and scientific enthusiasm, Prof. Massimo Battistini for his lessons on logic and writing, Prof. Elena Bacchelli for her suggestions on the use of gnomAD, and Prof. Stefano Alcaro who provided the computational resources required by the GBPM analysis. Finally, we thank Mr. George Wolf for the final proofreading the manuscript.

## Abbreviations

AA: amino acid
ACE2: Angiotensin-Converting Enzyme 2
COVID-19: Coronavirus Disease 2019
GBPM: Grid Based Pharmacophore Model
IEP: Interaction Energy point
MIFs: Molecular Interaction Fields
ORF: Open Reading Frame
PDB: Protein Data Bank
RBD: Spike Receptor Binding Domain with ACE2
RMSd: Root Mean Square deviation
SARS-CoV-2: Severe Acute Respiratory Syndrome Coronavirus 2

## Supplementary Files Description

Supplementary File 1: table of SARS-CoV-2 Spike mutations (source: GISAID database, 29 December 2020), indicating position, frequency in the sequenced SARS-CoV-2 genome and GBPM score (lower: predicted stronger effect in the Spike/ACE2 interaction).

Supplementary File 2: table of human ACE2 variants (source: gnomAD database, v3, 18 July 2020), indicating position, frequency in the sequenced SARS-CoV-2 genome and GBPM score (lower: predicted stronger effect in the Spike/ACE2 interaction).

Supplementary File 3: supplementary figures focusing on the most relevant mutations described in this study, with structural, chemical and positional considerations.

