## Supplementary File 3 for "Structural Genetics of circulating variants affecting the SARS-CoV-2 Spike / human ACE2 complex"

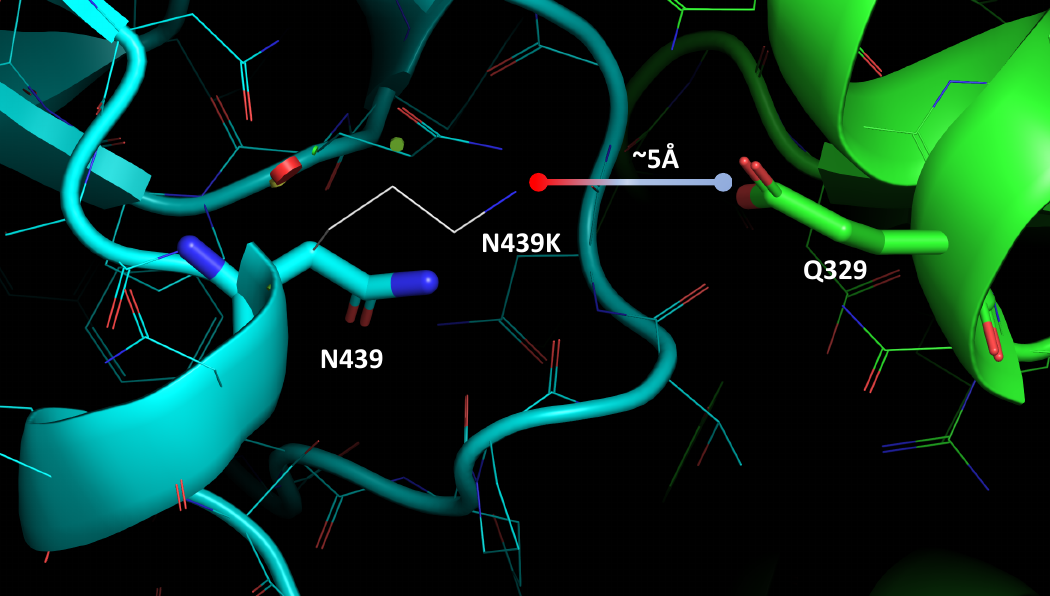

**Figure S1.** Spike N439K mutation. Spike and ACE2 proteins are depicted as cyan and green cartoon, respectively. Wildtype residues are reported as polytube, the mutation is depicted in white carbon wireframe.

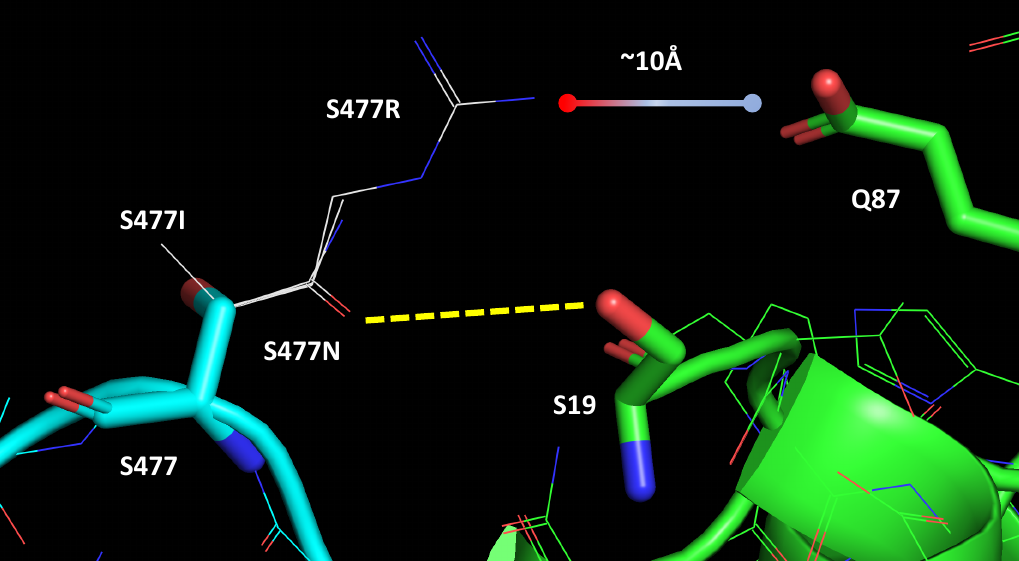

**Figure S2.** Spike S477N, S477I and S477R mutations. Spike and ACE2 proteins are depicted as cyan and green cartoon, respectively. Wildtype residues are reported as polytube, the mutations are depicted in white carbon wireframe. Yellow dotted line indicates hydrogen bond.

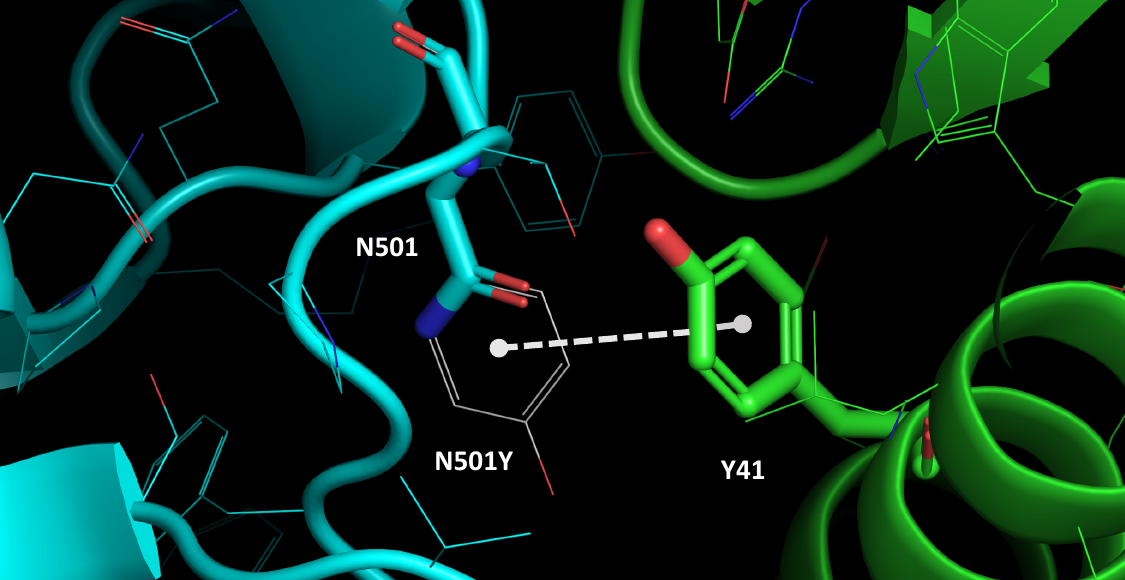

**Figure S3.** Spike N501Y mutation. Spike and ACE2 proteins are depicted as cyan and green cartoon, respectively. Wildtype residues are reported as polytube, the mutation is depicted in white carbon wireframe. White dotted line indicates pi-pi stacking interaction.

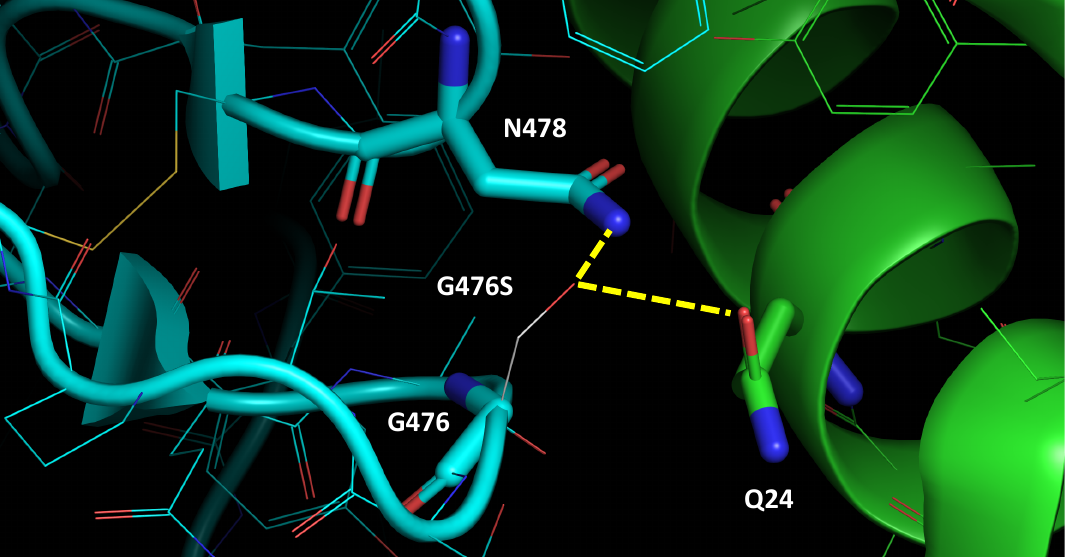

**Figure S4.** Spike G476S mutation. Spike and ACE2 proteins are depicted as cyan and green cartoon, respectively. Wildtype residues are reported as polytube, the mutation is depicted in white carbon wireframe. Yellow dotted line indicates hydrogen bond interaction.

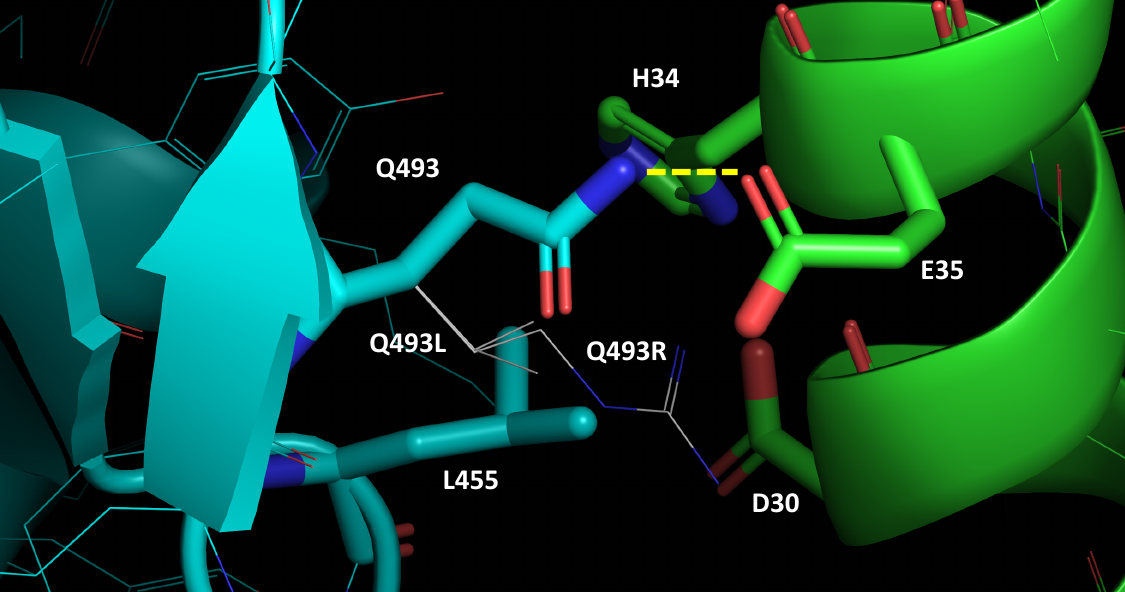

**Figure S5.** Spike Q493L and Q493R mutations. Spike and ACE2 proteins are depicted as cyan and green cartoon, respectively. Wildtype residues are reported as polytube, the mutations are depicted in white carbon wireframe. Yellow dotted line indicates hydrogen bond.

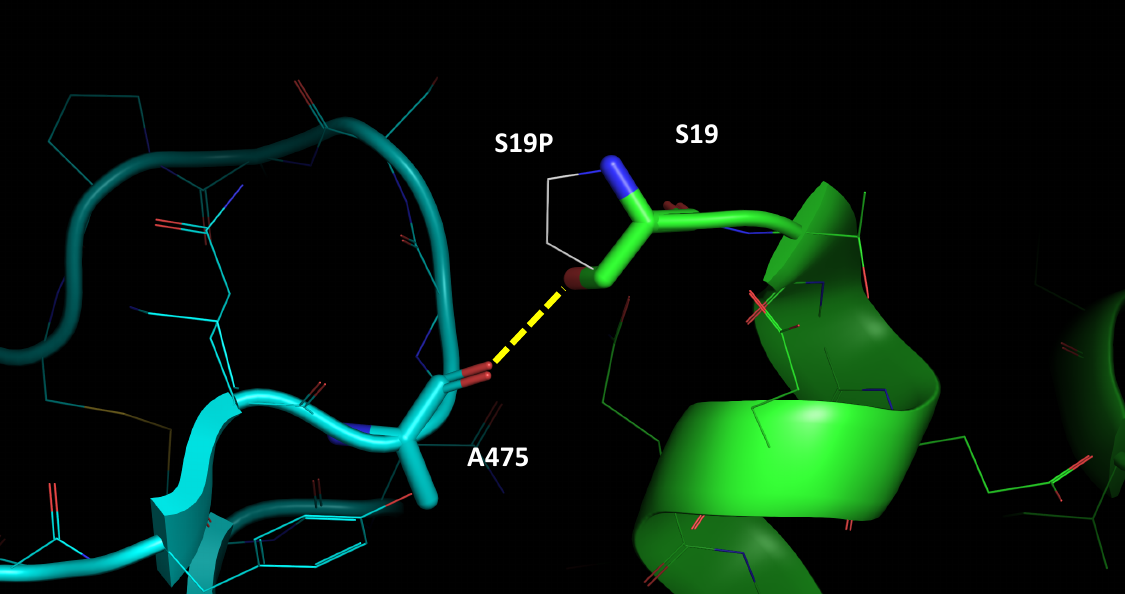

**Figure S6.** ACE-2 S19P mutation. Spike and ACE2 proteins are depicted as cyan and green cartoon, respectively. Wildtype residues are reported as polytube, the mutations are depicted in white carbon wireframe. Yellow dotted line indicates hydrogen bond.

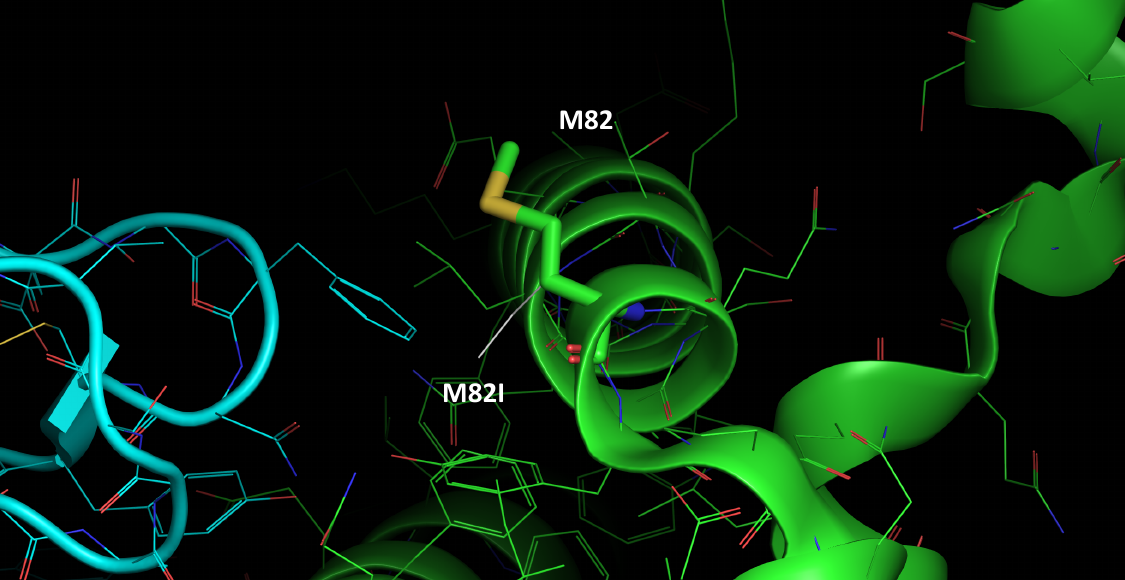

**Figure S7.** ACE-2 M82I mutation. Spike and ACE2 proteins are depicted as cyan and green cartoon, respectively. Wildtype residues are reported as polytube, the mutations are depicted in white carbon wireframe.

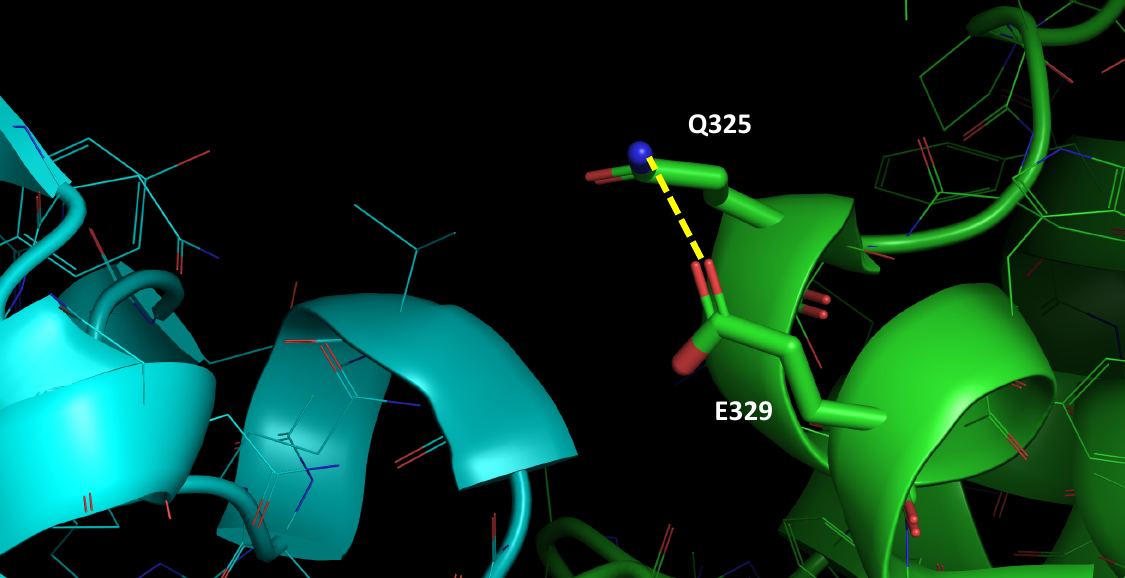

**Figure S8.** ACE-2 E329G mutation. Spike and ACE2 proteins are depicted as cyan and green cartoon, respectively. Wildtype residues are reported as polytube, the mutations is hidden. Yellow dotted line indicates hydrogen bond.
